# Dysregulated Alternative Splicing Landscape Identifies Intron Retention as a Hallmark and Spliceosome as a Therapeutic Vulnerability in Aggressive Prostate Cancer

**DOI:** 10.1101/634402

**Authors:** Dingxiao Zhang, Qiang Hu, Yibing Ji, Hsueh-Ping Chao, Amanda Tracz, Jason Kirk, Silvia Buonamici, Ping Zhu, Jianmin Wang, Song Liu, Dean G. Tang

## Abstract

Dysregulation of mRNA alternative splicing (AS) has been implicated in development and progression of hematological malignancies. Here we describe the first comprehensive AS landscape in the spectrum of human prostate cancer (PCa) development, progression and therapy resistance. We find that the severity of splicing dysregulation correlates with disease progression and establish intron retention (IR) as a hallmark of PCa stemness and aggressiveness. Systematic interrogation of 274 splicing-regulatory genes (SRGs) uncovers prevalent SRG mutations associated with, mainly, copy number variations leading to mis-expression of ~68% of SRGs during PCa evolution. Consequently, we identify many SRGs as prognostic markers associated with splicing disruption and patient outcome. Interestingly, androgen receptor (AR) controls a splicing program distinct from its transcriptional regulation. The spliceosome modulator, E7107, reverses cancer aggressiveness and abolishes the growth of castration-resistant PCa (CRPC) models. Altogether, we establish aberrant AS landscape caused by dysregulated SRGs as a novel therapeutic vulnerability for CRPC.

**Statement of significance:** We present the first comprehensive AS landscape during PCa evolution and link genomic and transcriptional alterations in SRGs to global splicing dysregulation. AR regulates splicing in pri-PCa and CRPC distinct from its transcriptional regulation. Intron retention is a hallmark for and spliceosome represents a therapeutic vulnerability in aggressive PCa.

## INTRODUCTION

Prostate cancer (PCa) still causes a significant mortality among men world-wide (1). The prostate is an exocrine gland containing mainly androgen receptor negative (AR^−^) basal and AR^+^ luminal epithelial cells, together with rare neuroendocrine (NE) cells (2,3). PCa predominantly displays a luminal phenotype and histologically presents as adenocarcinomas (Ad) largely devoid of basal cells (4). Most primary PCa (pri-PCa) are diagnosed as low to intermediate grade (i.e., Gleason grade ≤7), relatively indolent, and treated by radical prostatectomy and/or radiation with a good prognosis. Locally advanced (Gleason grade 9/10) and metastatic PCa are generally treated with androgen deprivation therapy (ADT) using LHRH agonists/antagonists, which block testicular androgen synthesis. Tumors that have failed this first-line therapy are termed castration-resistant PCa (CRPC) and further treated with anti-androgens such as enzalutamide (Enza) that interfere with the functions of AR. Enza extends CRPC patients’ lives by ~5 months but tumors inevitably become refractory to Enza. While the majority of CRPC and Enza-resistant tumors histologically present as adenocarcinoma (i.e., CRPC-Ad), a significant fraction (up to 25%) of them evolve to an aggressive, AR-indifferent disease with NE features called CRPC-NE (5). In general, all CRPC are relatively undifferentiated and, molecularly, basal/stem-like (6,7), highlighting lineage plasticity in facilitating treatment resistance and progression (8). Most metastatic CRPC (mCRPC), including both CRPC-Ad and CRPC-NE subtypes, remains lethal mainly due to our incomplete understanding of mechanisms underpinning CRPC emergence, maintenance and progression.

Dysregulation in pre-mRNA alternative splicing (AS) is emerging as a ‘hallmark’ of cancer (9,10). Nearly all multi-exon human genes undergo AS, a tightly regulated process that dramatically expands diversity of the transcriptome and proteome encoded by the genome (11). As an essential process for removing non-coding introns and ligating flanking exons to produce mature mRNA in eukaryotic cells, AS is performed by a dynamic and flexible macromolecular machine, the spliceosome. In addition to the core subunits that constitute five small nuclear ribonucleoprotein particles (snRNPs: U1, U2, U4, U5 and U6), the spliceosome contains many other auxiliary splicing-regulatory proteins (SRPs) including families of the serine- and arginine-rich (SR) proteins and heterogeneous nuclear ribonucleoproteins (hnRNP), and many other proteins that do not belong to these two families but have a role in modulating splicing (12). In this study, we refer to the genes encoding core subunits and the SRPs, broadly, as splicing-regulatory genes (SRGs) (Supplementary Table 1). Aberrant AS is prevalent in human cancers (13) and many cancer-specific splicing events contribute to disease development and progression (9,12). Since the initial discovery, via DNA sequencing, of frequent point mutations in the core spliceosome subunits in myelodysplastic syndromes and, later, in hematological malignancies (9), splicing dysregulation has been appreciated as a major contributor to cancer phenotypes. In parallel, therapeutic targeting of mis-splicing by small molecules presents a new approach for treating hematological malignancies bearing core subunit mutations (10,12) and solid tumors driven by MYC (14). Nevertheless, despite increasing elucidation of the global and cancer-associated splicing features by recent RNA sequencing (RNA-seq) analyses of primary tumors and normal tissues (13), the underlying molecular mechanisms as well as functional and clinical relevance of splicing misregulation in cancer, especially in solid tumors, remain largely undefined.

Importantly, many fewer recurrent mutations in the core spliceosome genes have been detected to date in solid tumors (15,16), suggesting a fundamental and mechanistic difference in splicing misregulation in hematological versus (vs.) solid cancers. Recently, global analyses of aberrant AS landscape across many human cancer types, including PCa, have been reported using RNA-seq data in TCGA (13,17–19), but these studies generally overlooked PCa and only analyzed pri-PCa, leaving behind life-threatening mCRPC. Furthermore, regarding the functional consequence of splicing dysregulation in PCa, previous studies have mainly focused on a few well-known genes typified by AR and CD44 (20), and the potential biological impact and clinical relevance of global splicing abnormalities in PCa remains unclear. Here, we specifically focus on PCa and provide, to our knowledge, the first comprehensive characterization of the AS landscape during disease development and progression and upon treatment failure. We report that intron retention (IR) represents the most salient and consistent feature across the spectrum of PCa entities and positively correlates with PCa stemness and aggressiveness. We also systematically analyze the dysregulated SRGs and correlate altered SRGs with aberrant AS patterns in PCa, and examine the deregulated pathways affected by aberrant splicing events. Finally, we demonstrate that splicing misregulation can be explored therapeutically for treating CRPC.

## RESULTS

### Increased AS Events Accompany PCa Development, Progression and Therapy Resistance

To determine the global dysregulation of AS in PCa development and progression, we employed two AS mapping algorithms, rMATS (21) and SUPPA (22), to annotate RNA-seq datasets encompassing pri-PCa and normal (N) prostate tissues (23), CRPC-Ad (24,25), CRPC-NE (5,26), and advanced PCa treated with hormonal therapy (26,27) (Fig. 1; Supplementary Fig. S1A). We defined ‘progression’ generally as stages beyond pri-PCa and as disease entities that were more aggressive in a comparative manner. Five main AS patterns, including alternative 3’ splice sites (A3), alternative 5’ splice sites (A5), mutually exclusive exons (MX), exon skipping (SE) and IR, were examined (Supplementary Fig. S1B). Splicing events with a cutoff of ΔPSI>0.1 and FDR<0.1 (for rMATS) or p<0.05 (for SUPPA) were considered statistically significant (see Methods).

**Figure 1.**
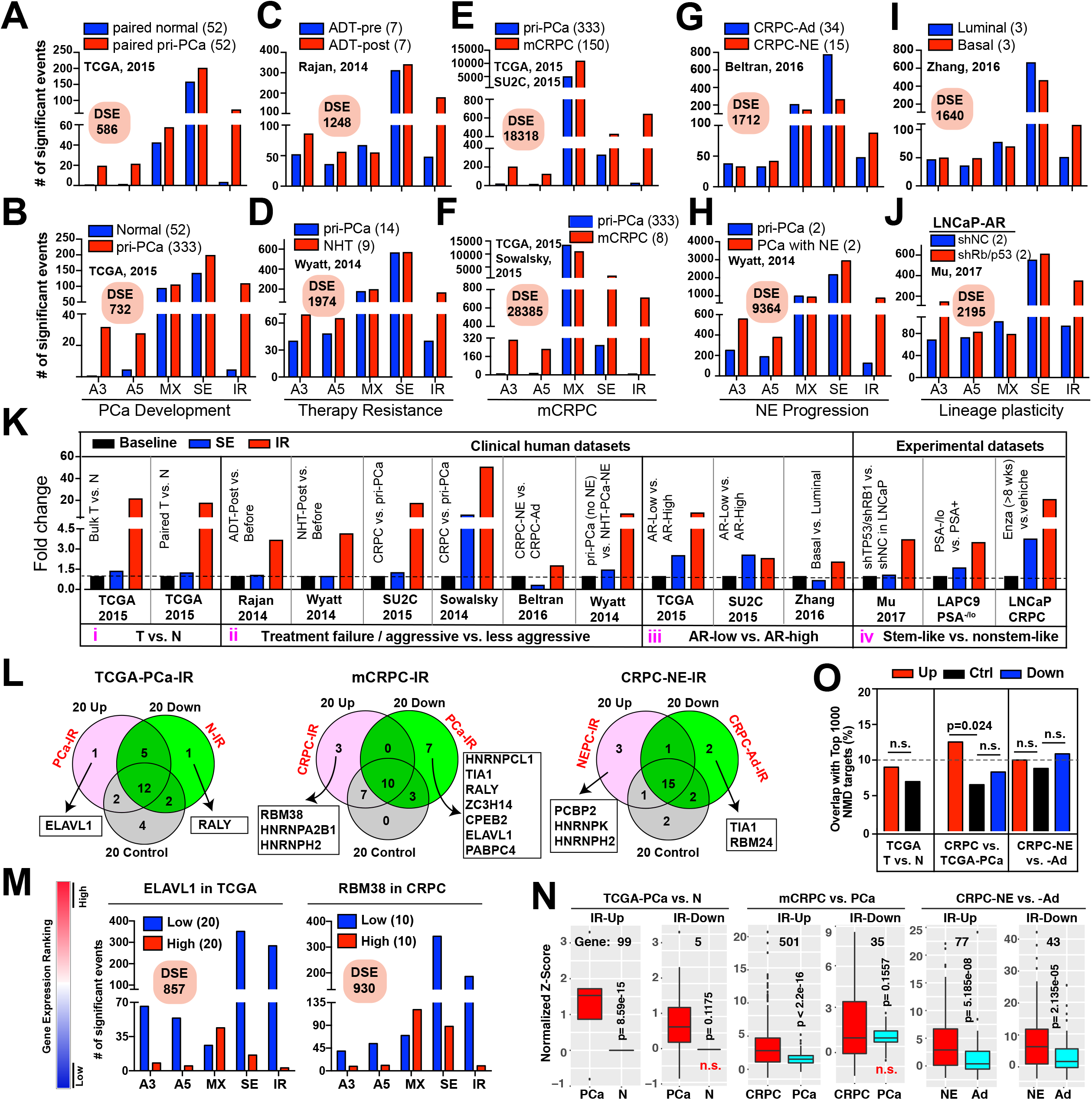
Splicing landscape identifies IR as a hallmark of PCa stemness and progression. (**A-J**) The landscape of AS during PCa development and progression. Two related datasets are interrogated for each PCa stage. Shown are splicing patterns and the number of DSEs decoded by rMATS. **(K)** Changes of IR and SE across the 14 comparisons detected by rMATS. “Baseline” refers to 1 as data presented as fold change in a comparative manner. **(L)** RBP motif analysis of retained introns specific to the PCa stages indicated. A total of 95 RBPs are examined and shown are the top 20 genes ranked by a binding score that takes into account both binding frequency and binding strength for each RBP. **(M)** DSEs associated with high or low expression level of ELAVL1 and RBM38 in pri-PCa and CRPC-Ad, respectively. **(N)** Pairwise comparison of expression of the genes showing significant IR events during PCa progression. Expression variability is quantified for each gene as a Z-score relative to the mean expression in normal prostate samples. Genes exhibiting both up-and down-regulated IR events are removed, and the resultant gene number is indicated. Significance was calculated by a paired Student’s *t*-test. **(O)** Overlap of significant IR-bearing genes with a high-confidence set of 1000 human NMD targets. Significance was calculated by a χ^2^ test. DSEs, differentially spliced events; RBP, RNA binding protein; NMD, nonsense-mediated mRNA decay.

Comparative analyses of either bulk or paired tumors and normal tissues indicated that pri-PCa possessed more AS events (~1.9 fold by rMATS; ~1.7 fold by SUPPA) with preferential increase in A3, A5 and IR (Fig. 1A and 1B; Supplementary Table S2). PCa post ADT (Fig. 1C) or subjected to neo-adjuvant hormone therapy (NHT; Fig. 1D) also displayed increased differentially spliced events (DSEs), suggesting a treatment-induced reshaping of global AS pattern that might have contributed to therapy resistance. Strikingly, mCRPC that had failed ADT and/or antiandrogens exhibited an exponential increase in DSEs, with noticeable increase in A3, A5, SE and IR (Fig. 1E and 1F). Within CRPC, CRPC-NE harbored a distinct splicing landscape relative to CRPC-Ad, although a notably smaller number of DSEs were observed than that in the CRPC-Ad vs. pri-PCa comparison (1712 vs. 18318; Fig. 1G). Interestingly, when comparing pri-PCa vs. pri-PCa with NE differentiation post NHT (26), we identified 9364 DSEs (Fig. 1H). Mapping with SUPPA revealed overall similar dysregulated AS patterns and progressive increase in DSEs in the spectrum of PCa development, therapy resistance, and progression (Supplementary Fig. S1C and S1D) albeit SUPPA by nature tended to detect more splicing events (see Methods). These results, taken together, suggest that PCa development is accompanied by increased AS events and that castration resistance and, in particular, metastasis, are characterized by further significant increases in AS events.

Since lineage plasticity facilitates therapeutic resistance and tumor progression (6,28), we determined the human prostate epithelial lineage-specific AS patterns as basal cells represent the main pool of prostate stem cells (SCs) and molecularly resemble aggressive PCa subtypes (7). Results revealed distinct AS profiles for prostatic basal vs. luminal cells, with more IR found in basal cells (Fig. 1I; Supplementary Fig. S1C). To determine whether basal-specific splicing profile also resembles that in aggressive PCa, we performed comparative gene set enrichment analysis (GSEA) and found that PCa with aggressive phenotypes (mCRPC and CRPC-NE) generally possessed a global basal-like AS profile (Supplementary Fig. S1E). Experimentally, silencing of tumor suppressors (TS) TP53 and RB1 in LNCaP/AR cells enables a lineage switch from AR^+^ luminal cells to AR^−^ basal-like cells (28). Consistently, a large number of DSEs were observed in LNCaP/AR cells with RB1/TP53 knockdown (Fig. 1J; Supplementary Fig. S1C), suggesting that plasticity driven by loss of RB1/TP53 is accompanied by a global shift in the AS landscape. Remarkably, GSEA indicated that the AS signatures of LNCaP/AR cells deficient in RB1/TP53 were significantly enriched in mCRPC compared to pri-PCa (Supplementary Fig. S1F). These results suggest that inherent lineage differences in normal prostate epithelial cells and induced lineage plasticity in PCa cells are also accompanied by dysregulated AS patterns that correlate with increased aggressiveness.

### AS Dysregulation Impacts PCa Biology

We explored the potential impact of AS dysregulation on PCa biology (Supplementary Fig. S2 and S3). By overlapping the splicing-affected genes (SAGs) and differentially expressed genes (DEGs), we observed only 9~20% of ‘overlapped’ genes (Supplementary Fig. S2A), suggesting that the majority of AS events minimally changes the bulk gene expression but may functionally tune transcriptomes (29). Gene ontology (GO) analysis (http://metascape.org) indicated that SAGs were enriched in many cancer-associated functional categories with both convergence (e.g., splicing, cell cycle and proliferation, cytoskeleton) and specificity identified at each PCa stage (Supplementary Fig. S2B-D). For instance, GO terms linked to ‘muscle and ion transport’, ‘lipid metabolism’, and ‘cell polarity’ were pri-PCa specific (Supplementary Fig. S2B) whereas GO terms ‘DNA damage’, ‘immunity’, and ‘nuclear pore’ were enriched in CRPC (Supplementary Fig. S2C), consistent with recent reports (30). Interestingly, and as expected, GO terms ‘SCs and development’ and ‘neuron and cell projection’ were greatly enriched in CRPC-NE (Supplementary Fig. S2D).

We further evaluated the potential functional consequences of AS dysregulation on PCa transcriptome by identifying transcript-level expression profiles using an isoform-specific alignment algorithm (31). As shown in Supplementary Fig. S3A, PCa at different stages exhibited distinct splice isoform signatures. For instance, the widely studied ARv7 was slightly upregulated in pri-PCa and, together with several other AR variants, was dramatically overexpressed in CRPC-Ad but not in CRPC-NE (due to loss of AR expression in NE tumors) (Supplementary Fig. S3B). CD44, a cancer stem cell (CSC) marker, plays versatile roles in metastasis with CD44-standard (CD44s) suppressing and CD44 variants (CD44v) promoting cancer cell colonization (32). Consistently, we observed a shift from no change in pri-PCa to a specific dysregulation of CD44 isoforms in mCRPC, with CD44s being downregulated in both CRPC-Ad and CRPC-NE and CD44v upregulated in CRPC-Ad (Supplementary Fig. S3C). The splicing program driving CRPC-NE emergence is scantly explored. Recently, an SE event leading to unique upregulation of a MEAF6 isoform containing exon 6 (i.e., MEAF6-204), but not the bulk mRNA, was reported in CRPC-NE (33). We observed similar results (Supplementary Fig. S3D), thus validating our analytic approaches. These analyses indicate that splicing abnormalities impact PCa biology by regulating cancer-related pathways, at least partially, via switching the isoform expression of key relevant genes.

### Increased IR as a Consistent Hallmark of PCa Progression, Stemness and Aggressiveness

Notably, we consistently observed increased IR across the spectrum of PCa development, progression and therapy resistance whereas the SE represented the most abundant splicing type (Fig. 1A-1J; Supplementary Table S2). We focused our subsequent studies on IR for it is the least studied AS type (10,34). We observed a >18 fold increase in IR in pri-PCa relative to normal tissues (Fig. 1K, i), consistent with a previous report that IR is common across multiple primary cancers (34). PCa progression is tightly associated with ADT failure and subsequent cellular plasticity towards stemness (35–37). In six different contexts, we consistently observed a preferential upregulation of IR in association with therapy-resistant, aggressive, and metastatic PCa (Fig. 1K, ii). Similar IR upregulation was observed in prostate tumors and epithelial cells displaying low vs. high AR activity (Fig. 1K, iii; see below). Interestingly, increased IR was also found in CSC-enriched PSA^−/lo^ cell population isolated from LAPC9 xenografts (38), basal-like LNCaP cells depleted of TP53 and RB1 (28), and LNCaP-CRPC cells that survived long-term Enza treatment (>8 weeks (39) (Fig. 1K, iv). Of note, SUPPA produced similar results (data not shown). These analyses link the upregulated IR with PCa stemness. We reanalyzed 3 recently published datasets that examined differentiation of different SC systems and also observed a positive correlation between IR and normal stemness (Supplementary Fig. S4A-C). Hence, in genetically matched hESCs – fibroblasts – iPS – fibroblasts system (40), ESCs lost IR during fibroblast differentiation while fibroblasts regained IR when they were reprogrammed to iPS cells (Supplementary Fig. S4A), in line with the earlier report (41). During spermatogenesis, spermatocytes displayed higher levels of IR than differentiated spermatids (Supplementary Fig. S4B), and these IR events were enriched in genes associated with gamete function (42). Finally, IR was found to be prevalent in stem-like, resting CD4^+^ T cells vs. functionally activated (differentiated) counterparts (Supplementary Fig. S4C), as reported previously (43).

We investigated the splicing ‘code’ of IR (44) in attempt to understand the molecular basis of preferential IR in aggressive PCa. Retained introns in normal tissues generally have weak 5’ and 3’ splice sites (44). Surprisingly, the splice site strength analysis did not reveal weak 5’ or 3’ sites in retained introns in pri-PCa, CRPC-Ad and CRPC-NE – in fact, CRPC-Ad showed stronger splice sites than pri-PCa (Supplementary Fig. S4D). Sequence feature analysis indicated that, compared with constitutive introns, IR in pri-PCa preferred introns with less GC content (GC%) and longer sequence length whereas CRPC-Ad specifically retained introns that were generally shorter without difference in GC% (Supplementary Fig. S4E). No feature variation was observed in the retained introns in CRPC-NE vs. CRPC-Ad (Supplementary Fig. S4E). Together, these results suggest that the prevalence of IR in PCa is not associated with weak 5’ and 3’ splice sites and may largely be *trans*-regulated.

To determine the potential *trans* factors (i.e., SRGs) that may preferentially regulate IR, we performed motif search for 95 RNA binding proteins (RBP) with known consensus motifs (45–47) on differentially splicing introns compared with constitutive introns. Based on an RBP-binding score for each factor, we chose top 20 genes for further analysis. As shown in Fig. 1L, we identified a few genes that may preferentially regulate IR for each specific comparison. Nonetheless, the majority of RBPs were shared by introns regardless of the IR status, suggesting that the spliceosome functions as a group rather than that one particular factor preferentially regulates one AS type. In support, we decoded the AS events associated with gene expression abundance by fractionating a cohort into two extremes (Fig. 1M). As expected, although the expression of ELAVL1 in pri-PCa and RBM38 in CRPC-Ad cohorts, respectively, both dramatically impacted IR, other splicing types were affected as well (Fig. 1M). Interestingly, ELAVL1 was not dysregulated in pri-PCa vs. normal tissues (FC=1.1). The discrepancy between a potential IR-inhibiting function of ELAVL1 and a marked increase in IR implied an involvement of other SRGs in preferential (or balanced) regulation of IR in pri-PCa. On the other hand, an IR-inhibiting function of RBM38 was consistent with its downregulation (FC=2.3) and an increase in IR in CRPC-Ad vs. pri-PCa (Fig. 1M). A TS role has been reported for RBM38 (48).

Subsequently, we interrogated potential biological impact of the upregulated IR on PCa biology. IR in normal conditions usually causes nonsense-mediated RNA decay (NMD) to downregulate gene expression (10,49). We compared the bulk RNA levels of IR-affected genes using two different mathematical methods and found that, surprisingly, these genes generally exhibited higher expression than their constitutively spliced counterparts (Fig. 1N; Supplementary Fig. S4F). To further strengthen our finding, we overlapped the IR-affected genes with a high-confidence set of human NMD targets (50) and found that only ~10% of genes in all groups were potentially targeted by NMD, although the genes with upregulated IR in CRPC tended to have slightly higher percentage (χ^2^ test, Fig. 1O). These results indicate that IR in PCa minimally causes NMD-mediated downregulation and these IR-bearing genes are thus likely functional. In support, GO analysis of IR-affected genes revealed that, in addition to commonly observed category of ‘splicing and RNA metabolism’, several distinct categories were enriched in aggressive PCa (Supplementary Fig. S4G-I). For example, GO terms ‘stress response’, ‘DNA repair’ and ‘cancer-related signaling’ (e.g., ERBB, NOTCH, WNT) were unique to CRPC-Ad (Supplementary Fig. S4H) whereas ‘hormone transport’ and ‘SC & development’ were strongly associated with androgen-insensitive and CSC-enriched CRPC-NE (Supplementary Fig. S4I).

### AR Regulates a Splicing Program Distinct from the AR-Regulated Transcriptome

AR is obligatory for pri-PCa growth and continues to be expressed and functionally important in CRPC (51). ADT promotes a stem-like phenotype in PCa (52) and relapsed tumors often exhibit enhanced SC properties (8,38,53). We set out to determine whether AR may drive splicing dysregulation seen in PCa development and progression. We first established an AR activity score based on the Z-scores calculated from the expression of 20 experimentally validated AR targets (23). The TCGA cohort bearing ‘uninterrupted’ intrinsic AR heterogeneity (23) and CRPC-Ad cohort bearing ‘twisted’ AR activity by treatments (24) were then fractionated into high and low AR activity groups, followed by splicing analyses. Not surprisingly, primary tumors with low vs. high AR activities displayed a significant difference in AS landscape (Fig. 2A), and this difference was amplified in CRPC (Fig. 2B), implicating AR signaling in modulating global AS. Of note, we observed no association between AR genomic alterations and its potential splicing-modulating activity (Fig. 2C) since AR is rarely altered in pri-PCa but frequently amplified in mCRPC (24). To assess the impact of AR-associated splicing on AR-regulated gene expression, we compared the SAGs with DEGs identified in the AR-low vs. AR-high comparisons. Surprisingly, only 2% of SAGs overlapped with the DEGs in pri-PCa, although this overlap was increased to 23.2% in CRPC-Ad (probably due to a much-enlarged repertoire of AR-regulated molecular events in CRPC) (Fig. 2D). Thus, AR activity-associated AS events exerted a limited impact on the AR transcriptional targets, leading us to hypothesize that AR regulates a splicing program distinct from its transcriptional regulation. In support, when we extended the comparison to three other well-defined AR-target gene sets ((54), and two in this study, see below), we observed generally <4% overlaps across all comparisons (Fig. 2E).

**Figure 2.**
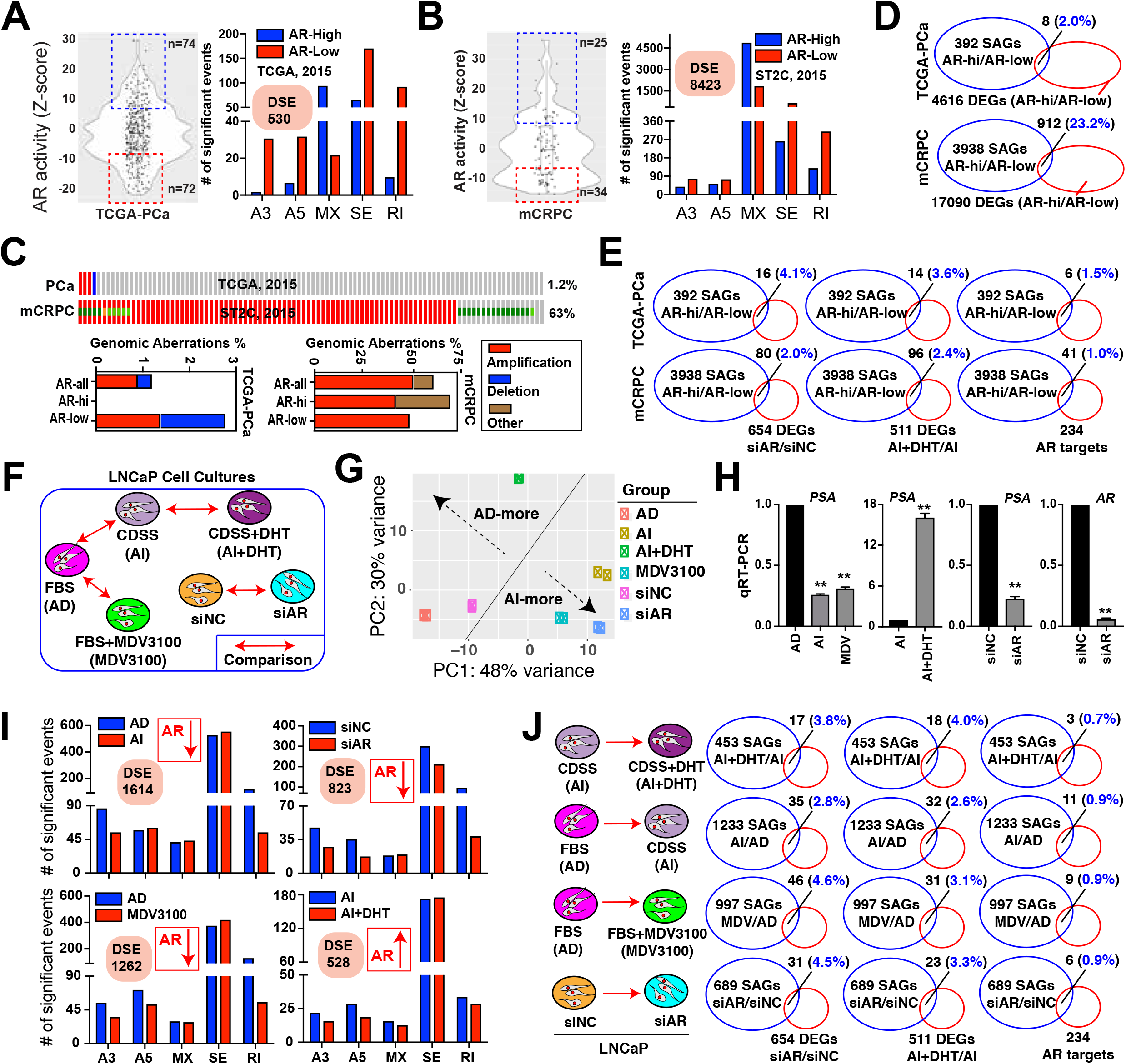
AR activity impacts AS landscape distinctively from its regulation of transcription. (**A-B**) DSEs associated with high and low AR activity (cutoff, Z-score >7 or <-7) in pri-PCa (A) and CRPC-Ad (B), respectively. AR activity (see Methods) was used to fractionate patients cohorts followed by splicing analysis by rMATS. (**C**) Genomic alterations do not contribute to the diversity of AR activities across PCa populations. Shown are frequency and AR mutation types observed in TCGA and CRPC cohorts. AR activities of samples grouped as in A were displayed. (**D-E**) Overlap between SAGs and DEGs (D) and between SAGs and three sets of AR-regulated genes (E) in indicated contexts. The number in parentheses denotes a percentage of overlapped genes proportioned to all SAGs. Circles are not drawn to scale. (**F-H**) Experimental design (F), principal component analysis (PCA) showing proper clustering of samples (G), and qPCR validation of intended modulations of AR signaling in LNCaP cells (H). (**I**)The AR-regulated AS program in PCa cells. Shown are the DSEs associated with high (up arrow) or low (down arrow) AR activity in LNCaP cells detected by rMATS. (**J**)Overlap between SAGs and three sets of AR-regulated genes in indicated contexts. The number in parentheses denotes a percentage of overlapped genes proportioned to all SAGs. Circles are not drawn to scale. DSEs, differentially spliced events; DEGs, differentially expressed genes; SAGs, splicing-affected genes.

To experimentally validate our hypothesis, we treated AR^+^ LNCaP cultures with various regimens to modulate AR activity (Fig. 2F). Cells cultured in regular fetal bovine serum (FBS)-containing medium represent an androgen-dependent (AD, or androgen-sensitive) state. Cells grown for 4 days in medium containing charcoal/dextran stripped serum (CDSS) or treated with Enza (i.e., MDV3100, 10 μM) were considered androgen-independent (AI). We also utilized siRNA to genetically silence AR. Finally, cells primed with CDSS for 3 days were treated with 10 nM dihydrotestosterone (DHT) for 8 h to restore AR signaling. Deep RNA-seq was performed in biological duplicates on abovementioned LNCaP cultures (Fig. 2F; Supplementary Fig. S5A). Principal component analysis (PCA) indicated that samples were properly clustered (Fig. 2G) and AR signaling was effectively modulated as intended, evidenced by expression levels of AR and PSA and by GSEA of AR gene signature (Supplementary Fig. S5B) and by quantitative reverse transcription PCR (qRT-PCR) validation (Fig. 2H). Pairwise comparisons uncovered significant differences in DSEs in cells exhibiting high vs. low AR activities (Fig. 2I; Supplementary Fig. S5C). Also, reanalysis of a recent RNA-seq dataset (GSE71797) (55) confirmed that in response to R1881 (24~48 h), activated AR signaling reshaped the AS landscape in 3 AR^+^ PCa cell models (i.e., LNCaP, VCaP, 22Rv1) (Supplementary Fig. S5D). Similarly, by categorizing the DEGs identified in cells depleted of AR (siAR vs. siNC) or treated with DHT as AR-target sets, we found that, strikingly, these two sets, together with a previously reported AR signature (54), minimally overlapped with the SAGs (<5%) defined in all different contexts (Fig. 2J; Supplementary Fig. S5E). Collectively, we conclude that AR regulates a set of AS-bearing genes distinct from its transcriptional targets, with or without the presence of androgen.

We also investigated whether AR might specifically regulate IR, as tumors and basal cells with low canonical AR activity were associated with increased level of IR (Fig. 1K). Surprisingly, our work in LNCaP system revealed that a decrease in AR activity resulted in no increase, but a decrease, in IR while stimulation of AR-mediated transcription failed to appreciably repress IR (Supplementary Fig. S5F). To further test this experimentally, we utilized a quantitative reporter system (56) in which a 132-nucleotide chimeric β-globin/immunoglobulin intron was inserted into the firefly luciferase gene (Supplementary Fig. S5G). Dual luciferase assays (Supplementary Fig. S5G) indicated that consistent with previous reports (56), splicing conferred an advantage to gene expression in that equal amounts of transfected plasmids generated higher signals from intron-containing than intronless luciferases (Supplementary Fig. S5H, left). However, luciferases with or without intron generated a similar pattern of signal changes across conditions with dampened or enhanced AR signaling (Supplementary Fig. S5H, right), suggesting that AR does not specifically regulate IR in AR^+^ PCa cells.

### Distinct Genomic Alterations in SRGs Impact AS and Associate with PCa Aggressiveness

Recent genomic sequencing efforts have revealed the global mutational landscapes of PCa during development and progression (5,23,24,51,57–61), almost all of which focused their initial analysis on known PCa-related genes and pathways (e.g., AR, PTEN/PI3K, TP53, RB1, DNA repair, ETS fusion) whereas alterations in SRGs were overlooked due to a low mutation frequency at individual gene level. Moreover, point mutations in spliceosome core genes have been recognized as a key driver in hematological cancers (12). We explored the molecular mechanisms underpinning the AS dysregulation in PCa by compiling and curating a catalog of 274 SRGs (Supplementary Table S1) and systematically surveying their mutational landscape (Fig. 3; Supplementary Fig. S6-S8). We interrogated 11 available large-scale clinical datasets in cBioportal (62) and excluded 3 from further analysis due to limited information available (Supplementary Table S1). The remaining 8 were categorized as pri-PCa and CRPC datasets. Fig. 3A and 3B showed the mutational landscape of top 15 altered SRGs in representative pri-PCa and CRPC datasets (also see Supplementary Fig. S6 and S7), respectively. Based on this global mutational landscape, several interesting patterns emerged. **First**, genomic deletions of SRGs in pri-PCa and amplifications of SRGs in CRPC represented the most prevalent forms of alterations, among others (Fig. 3C and 3D). **Second**, the frequently deleted and amplified genes often co-occurred with the deletion of TS genes and amplification of oncogenes, respectively (also see Supplementary Fig. S8A). For example, ENOX1, WBP4, HNRNPA1L2 and RB1 were co-localized and co-deleted on Chr13q (p=5.16E-42; Supplementary Table S1). On the other hand, KHDRBS3, PABPC1, ESRP1, PUF60 were co-amplified with MYC on 8q (p≤1.50E-15). **Third**, most of the SRGs mutated at low frequency, as only 20 (7.3%) and 29 (10.58%) of the 274 SRGs were mutated at a rate of ≥5% in TCGA-PCa and SU2C-CRPC cohorts, respectively. Consequently, the mutation burden in sum is predominantly contributed by the top 20 altered genes (Fig. 3E; Supplementary Table S1). **Fourth**, chromosomal distribution of mutated SRGs (≥5%) revealed that, except the top altered genes, the majority of SRGs were localized outside the previously reported hotspots (23,61) (Supplementary Fig. S8B), in line with their low mutation rates. In aggregate, our data indicate that, albeit a low alteration frequency at individual gene level, SRGs, collectively, represent a frequently mutated pathway in PCa, as ~31-68% and 87-94% of patients with pri-PCa and CRPC, respectively, harbor at least one mutation of one SRG (Supplementary Table S1).

**Figure 3.**
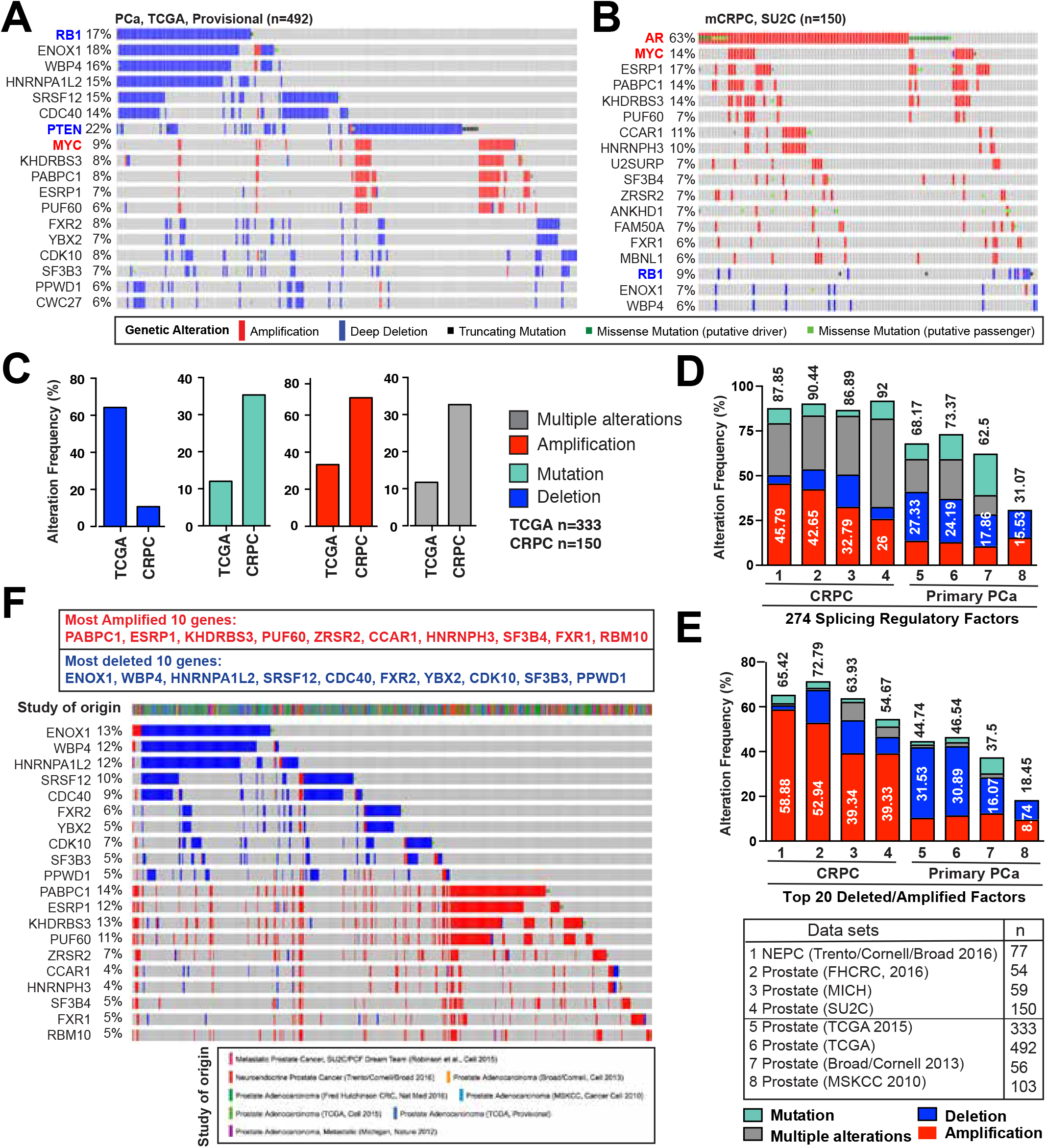
The mutational landscape of SRGs in human PCa. (**A-C**) A comprehensive survey for genomic alterations in 274 SRGs in available clinical cohorts in cBioportal. The top 15 mutated SRGs are shown in the representative pri-PCa (A) and metastatic CRPC-Ad (B) cohorts. Frequently deleted RB1 and PTEN (colored in blue) and amplified MYC and AR (colored in red) are included as reference genes. Each bar represents the alteration status of an individual gene for a single patient and the percentage of alterations for each gene in the indicated cohort is provided. Shown in C are bar graphs summarizing the cumulative genomic alterations of SRGs in the largest and representative pri-PCa (TCGA) and CRPC (SU2C) cohorts. (**D-E**) Bar plots illustrating the cumulative aberration frequencies of all 274 SRGs combined (D) and the top 20 mutated SRGs (10 most amplified and 10 most deleted) (E) across all cohorts, with numbers above and within the bars representing the total frequency and a frequency of amplification or deletion of indicated genes, respectively. (**F**) Integrated mutational landscape of top 20 mutated SRGs in PCa showing mutual exclusivity, in large part, between deletions and amplifications of SRGs. See Table S1 for detail.

Evolutionarily, deletion and amplification of selective SRGs might represent early and late events, respectively, in PCa pathogenesis (Fig. 3C). In support, group analysis of top altered SRGs showed that deletion of SRGs did not, whereas amplification of SRGs did, associate with increased Gleason grade (not shown), highlighting a potential survival advantage of clones harboring SRG amplifications over deletions during PCa progression. This notion is further supported by a recent study showing that focal genomic amplifications represent a rapid adaptation to selection pressure and a driving force in metastatic CRPC (63). We also observed an overall increase (e.g., 8% to 14% for KHDRBS3, 7% to 17% for ESRP1) and decrease (e,g., 18% to 7% for ENOX1, 16% to 6% for WBP4) in the frequencies of amplified and deleted genes, respectively, in CRPC vs. pri-PCa (Fig. 3A and 3B). Interestingly, SRG deletions and amplifications seemed to be mutually exclusive (Fig. 3F).

We reasoned that copy number variation (CNVs) in SRGs might lead to their differential mRNA expression, which in turn might be tied to splicing misregulation in PCa. Indeed, gene expression analysis for top altered SRGs in both pri-PCa and CRPC indicated that deletion and amplification generally correlated with loss and gain of mRNA expression, respectively (Supplementary Fig. S9A-B). Oncomine Concept analysis revealed 72 and 74 dysregulated SRGs (p<0.05) in pri-PCa and CRPC, respectively (Supplementary Fig. S9C). In RNA-seq datasets, 33, 89 and 45 SRGs were significantly deregulated in pri-PCa (vs. normal tissues), CRPC-Ad (vs. pri-PCa) and CRPC-NEPC (vs. CRPC-Ad), respectively (Supplementary Fig. S9D). Furthermore, an RNA-seq examining the response of advanced PCa to ADT (27) revealed 19 DEGs, and, of interest, an exclusive overexpression of 7 genes was identified in basal vs. luminal cells (Supplementary Fig. S9D). Notably, many of the top amplified and deleted SRGs were also found to be, correspondingly, overexpressed or downregulated in PCa at the population level (Supplementary Fig. S9E). An integrated summary (Fig. 4; Supplementary Table S3) revealed that, in total, 186 out of 274 (67.9%) SRGs were mis-expressed at different stages of PCa, with more dysregulated SRGs found in CRPC, implicating a potential dependency of aggressive PCa on spliceosome activity.

**Figure 4.**
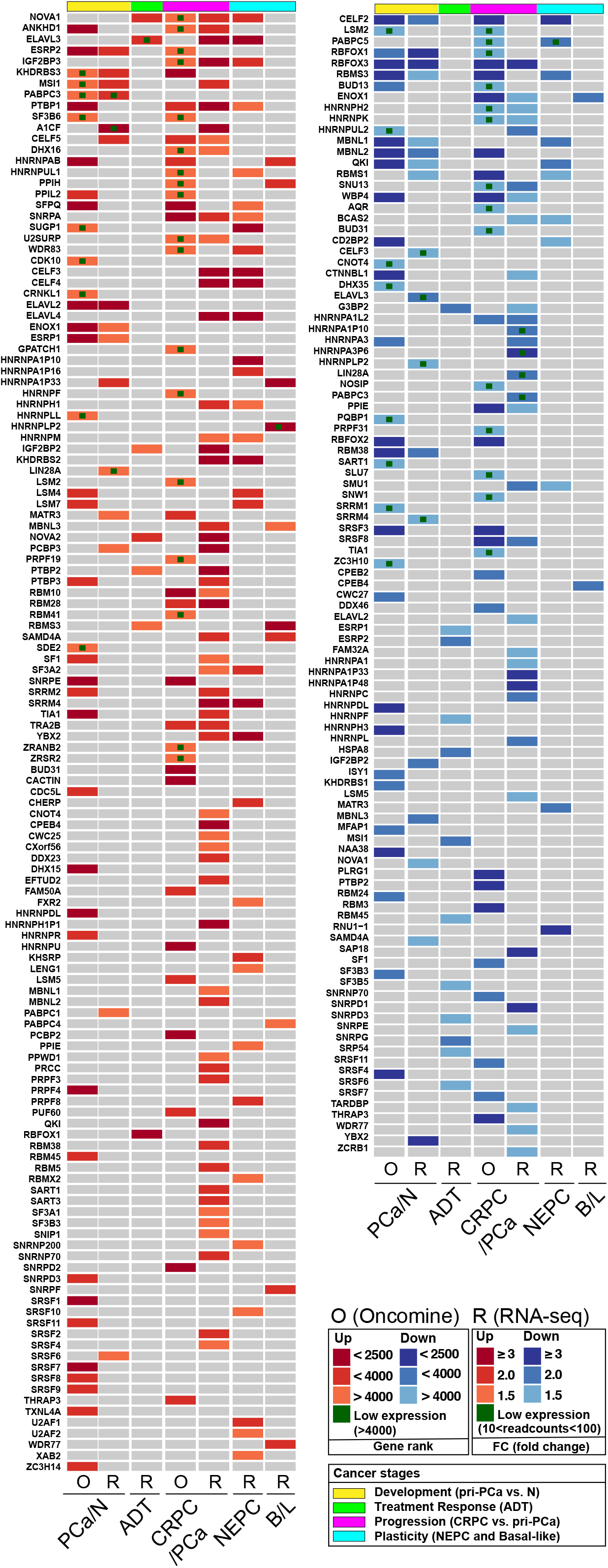
Dysregulation of SRGs in PCa. Integrated heatmap of differentially expressed SRGs identified in Oncomine (p<0.05) and RNA-seq (fold-change (FC)≥ 1.5 and FDR<0.1). In Oncomine (O), the medium-rank of <2500, <4000, and >4000 for a gene denotes high, moderate and low level of expression, respectively. For visualization, DEGs revealed by RNA-seq (R) data are categorized into three groups according to FC differences (FC≥ 3, ≥ 2 and ≥ 1.5). Based on pairwise comparisons, the stages of PCa are defined as tumor development (pri-PCa vs. normal tissues), ADT treatment response (ADT-after vs. -before), CRPC progression (CRPC-Ad vs. pri-PCa) and plasticity (CRPC-NE vs. CRPC-Ad).

To further explore the clinical relevance of SRGs, we assessed the prognostic values of dysregulated SRGs in patient’s outcome. We systematically surveyed the 186 misregulated SRGs in 7 Oncomine datasets containing patient survival information and identified two types of ‘prognostic’ SRGs: unfavorable genes whose higher expression correlated with poor patient survival and favorable genes whose higher expression correlated with better patient survival (Fig. 5A; Supplementary Table S4). In general, we observed a consistency between overexpressed SRGs and unfavorable prognostic genes, but not for down-regulated genes and favorable prognostic genes (Supplementary Table S4). Interestingly, although different datasets revealed varying numbers of prognostic genes (Supplementary Table S4), we identified more SRG genes classified as unfavorable genes (Fig. 5B). Together with the mutational landscape (Fig. 3) and deregulated expression patterns of SRGs (Fig. 4) that cooperatively indicated a potential dependency of CRPC on spliceosome activity, this would strongly suggest that SRGs, mostly, play oncogenic roles in PCa progression. Importantly, most of the identified prognostic genes have not previously been linked to PCa patient survival. Towards a better use of these prognostic SRGs in heterogeneous PCa, we established two gene signatures based on the consistency of the survival results seen in the 7 datasets, corresponding to unfavorable signature (13 genes, SRSF1, KHDRBS3, ESRP1, HNRNPH1, U2SURP, LSM5, TIA1, CHERP, HNRNPR, HNRNPH2, HNRNPH3, HNRNPAB and KHDRBS1, with each showing consistent unfavorable prognosis in ≥3 datasets) and favorable signature (13 genes, MFAP1, SF3A2, GPATCH1, XAB2, CELF2, SF3A1, SAP18, SRP54, PPIL2, SF1, MATR3, ELAVL4 and CDK10 with each showing consistent favorable prognosis in ≥2 datasets). We found that patients whose cancer gene expression enriched for the unfavorable or favorable signatures had a worse or a better survival outcome, respectively (Fig. 5C), suggesting a utility of SRGs as prognostic biomarkers. To further study underlying link between prognostic SRGs and splicing dysregulation, we investigated impact of unfavorable signature on disease aggressiveness and splicing in TCGA cohort. As expected, the unfavorable signature score positively and negatively associated with the tumor grade and disease recurrence, respectively (Fig. 5D and 5E). Importantly, primary tumors expressing highly or lowly the unfavorable signature exhibited distinct splicing landscapes, with total DSEs (1.73 fold) and IR (18.91 fold) being specifically upregulated in the high group (Fig. 5F).

**Figure 5.**
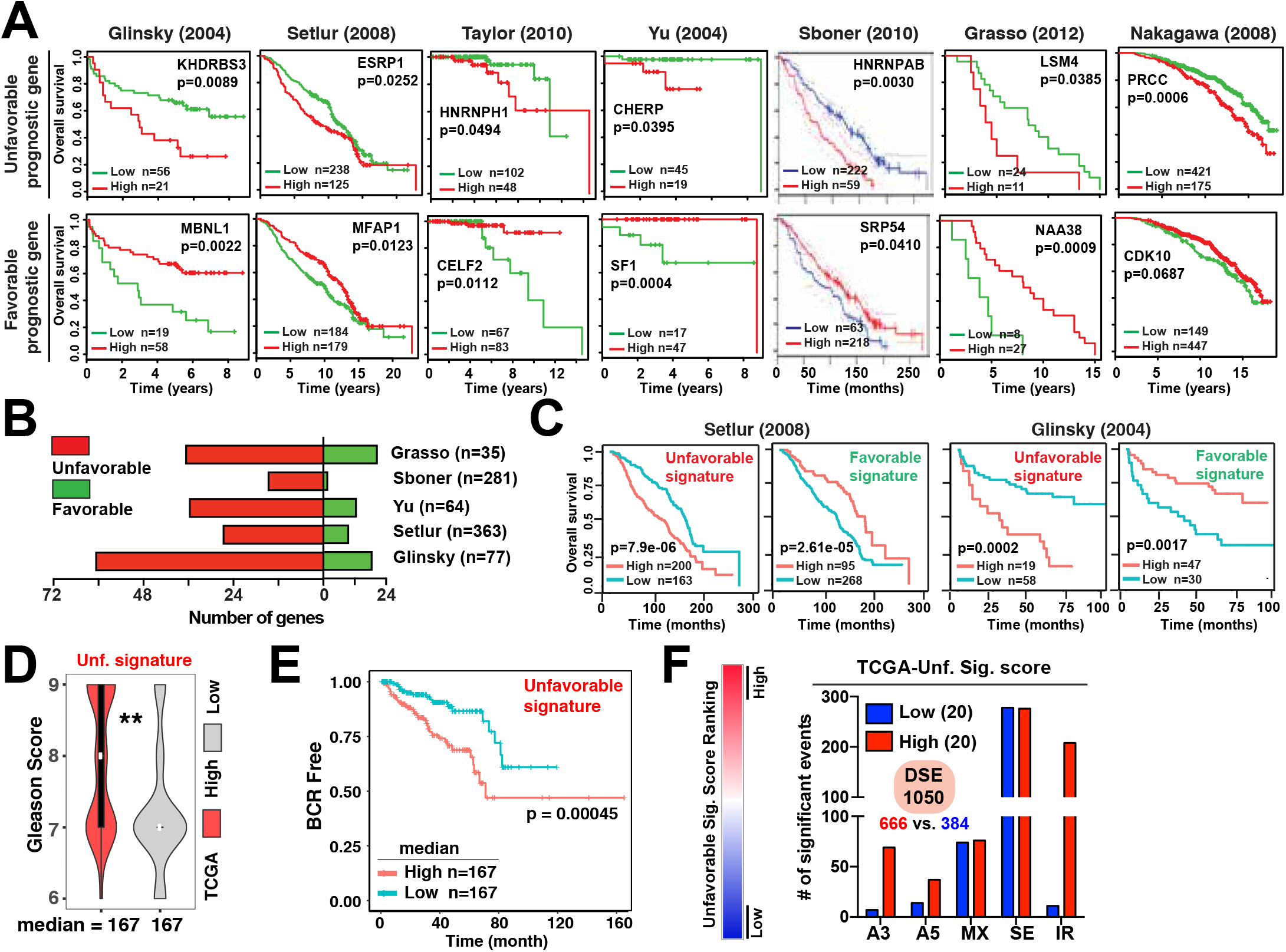
SRGs are prognostic and associated with splicing dysregulation. (**A**) Examples of Kaplan-Meier plots for unfavorable and favorable genes associated with patient overall survival in 7 different cohorts. (**B**) Numbers of genes showing favorable and unfavorable prognostic effects in 5 distinct cohorts. Patient numbers for each cohort are shown in parentheses. (**C**) Meta-analysis showing higher level of unfavorable signature and lower level of favorable signature correlating with reduced overall patient survival, respectively. Data were based on the Setlur and Glinsky studies. (**D-E**) Unfavorable signature is associated with higher Gleason score (D) and higher level of unfavorable signature positively correlates with disease recurrence in TCGA cohort (E). (**F**) DSEs associated with high or low expression level of unfavorable signature in primary PCa cohort. The p value was calculated using Student’s *t*-test (D) and log-rank test (A, C, E) *p< 0.05. See Table S4 for details.

### CPPC Cells Are Sensitive to Pharmacological Spliceosome Inhibition In vitro

Based on the observations that mCRPC possess frequent amplifications in (Fig. 3) and deregulation of SRGs (Fig. 4), that higher expression of SRGs predicts worse outcome (Fig. 5), and that modulation of AR activity reshapes PCa-associated AS landscape (Fig. 2), we hypothesized that *spliceosome may represent a preferential CRPC dependency*, thus offering a therapeutic opportunity. To test this, we first analyzed the mutational profiles of SRGs in 7 PCa cell lines with increasing aggressiveness and found that the AR^+^ and relatively indolent PCa cells tended to have more SRG deletions whereas AR^−^ and aggressive PCa cells showed more SRG amplifications (Supplementary Fig. S10A). In particular, LNCaP and PC3 cells resembled pri-PCa and CRPC, respectively, with respect to SRG mutation profiles (Supplementary Fig. S10A). We retrieved two large-scale RNAi screening data (Novartis Project Drive (64) and Broad Project Achilles (65)) and performed GSEA on ranked lists of essential genes. We observed that aggressive AR^−^ PCa cell lines exhibited a preferential enrichment on two splicing pathway signatures (Supplementary Fig. S10B). By contrast, AR signaling and MYC signatures were enriched in AR^+^ LNCaP and 22RV1 vs. AR^−^ DU145 cells, respectively (Supplementary Fig. S10C). These analyses support the postulate that AR^−^, androgen-independent PCa cells may be particularly dependent on the spliceosome activity.

We subsequently tested this postulate using spliceosome inhibitors. Several microbial products, including Pladienolide B and its derivative E7107 have been shown to bind and inhibit the SF3B1 complex and manifest anti-cancer activities (12,20). The E7107 compound represented the first-in-class spliceosome inhibitor that underwent phase I clinical trial (66). We found that PCa cells exhibited preferential sensitivity to E7107 relative to non-tumorigenic prostate epithelial cells RWPE1, with PC3 being more sensitive than LNCaP cells (Fig. 6A; Supplementary Fig. S10D). Experiments with Pladienolide B confirmed PC3 as the most sensitive line (Fig. 6B). While a long-term E7107 treatment (6~7 days) induced massive cell death (Fig. 6A; Supplementary Fig. S10D), shorter (<3 days) treatments generally elicited limited apoptosis but instead arrested PCa cells at the G2/M phase of the cell cycle (Fig. 6C). Treatment of PCa cells with E7107 for 20~48 h also inhibited cell migration and invasion, as measured by both Boyden chamber (Fig. 6D; Supplementary Fig. S10E) and scratch-wound (Supplementary Fig. S10F) assays. Importantly, treatment of PCa cells with 5 nM E7107 for 6 h dramatically reshaped the splicing pattern of the selected genes (Supplementary Fig. S10G; also see below), suggesting an on-target effect of the drug.

**Figure 6.**
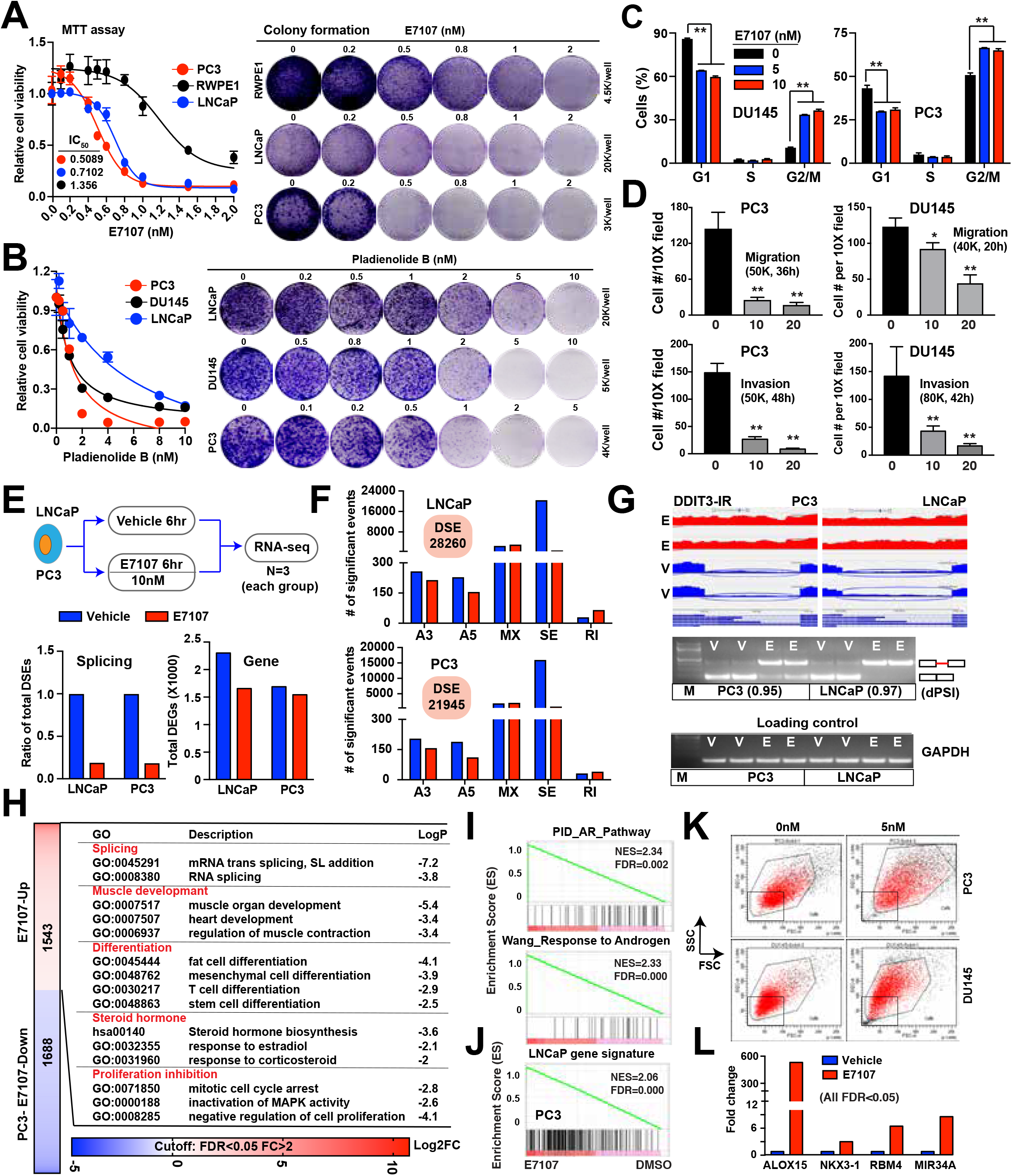
CRPC cells are sensitive to pharmacologic modulation of spliceosome activity. (**A-B**) Cell proliferation (MTT; left) and colony formation (right) assays in indicated cell lines treated with varying concentration of E7107 (A) or Pladienolide B (B) *in vitro*. (**C-D**) Cell cycle analysis (C) and migration and invasion assays (D) in indicated PCa cells treated with varying concentrations of E7107. Results shown were representative of 2–3 repeat experiments. For D, data represent mean±S.D. from cell number counting of 5–6 random high magnification (X20) areas. The P value was calculated using Student’s *t*-test. *P<0.05 and **P<0.01. (**E**) Effect of 10nM E7107 on PCa transcriptome *in vitro*. Shown are schematic of RNA-seq experiments (top) and the number of total DSEs (bottom left) and DEGs (bottom right) identified upon E7107 treatment in indicated PCa cells. (**F**) AS pattern showing that E7107 reshapes the splicing landscape of PCa cells indicated. (**G**) Sashimi plots visualization and RT-PCR validation of IR in DDIT3 gene after an acute E7107 (10 nM, 6 h) treatment. The event ΔPSI values calculated by rMATS were provided in parentheses. (**H**) GO analysis of genes upregulated at bulk RNA level in PC3 cells after E7107 treatment (10nM, 6 h). (**I-J**) GSEA showing enrichment of AR signaling related signatures (I) in E7107-treated PC3 cells, indicating that E7107 reprograms the AR^−^ PC3 cells back into relatively AR^+^ LNCaP-like cells. In support, a LNCaP gene signature (defined as top 300 genes solely expressed or overexpressed in LNCaP compared with PC3) was significantly enriched in PC3 cells treated with E7107 (J). (**K**)Representative FACS plots of PC3 and DU145 cells treated with E7107 (5 nM) for 3 days showing an increase in cell size. (**L**)Upregulation of tumor suppressors (ALOX15, KNX3-1, RBM4 and MIR34A) in PC3 cells after E7107 treatment.

### E7107 Molecularly ‘Reverses’ PCa Cell Aggressiveness by Inhibiting Spliceosome Activity

To uncover the mechanisms of action of E7107 in PCa, we treated LNCaP and PC3 cells with the drug for 6 h followed by deep RNA-seq (Fig. 6E). No gross defects were observed in cell growth (Supplementary Fig. S11A), but, as expected, E7107 dramatically inhibited the AS globally in both cell types (Fig. 6E) with SE being the major splicing type affected (Fig. 6F). Sashimi plot visualization of the sequencing data and RT-PCR analysis validated splicing analysis (Fig. 6G; Supplementary Fig. S11B). We performed GO analysis on genes showing down-regulated splicing events. Analysis of the top 1000 genes with significant SE events inhibited by E7107 in PC3 cells revealed that many GO terms associated with cancer-promoting functions, e.g., cell cycle and proliferation, DNA repair, splicing, and cancer pathways, were markedly enriched (Supplementary Fig. S11C; Supplementary Table S5), suggesting that E7107 inhibits splicing of a subset of PCa-associated genes important for survival. At the gene expression level, E7107 reshaped the transcriptomes and exhibited a slightly suppressive effect, especially in LNCaP cells, on transcription (Fig. 6E; Supplementary Table S6). qRT-PCR analysis validated DEGs identified in RNA-seq (Supplementary Fig. S11D).

We also performed GO analysis of DEGs upregulated after E7107 treatment. In AR^+^p53^+^ LNCaP cells, four main categories of related GO terms were identified (Supplementary Fig. S11E) with ‘Splicing’ being the most significant one, consistent with a recent report (67). AR and its target expression and AR signaling were not significantly affected by E7107 in LNCaP cells (Supplementary Fig. S11F). Interestingly, p53 was activated, along with several other TS genes including RBM4 (68) and MIR34A (69) (Supplementary Fig. S11E and S11G). Consistently, GO terms ‘cell cycle arrest’ and ‘differentiation’ were enriched (Supplementary Fig. S11E). We have previously shown that the LNCaP gene expression profile resembles that in pri-PCa (7). GSEA of gene signatures specific to normal prostate tissues vs. pri-PCa revealed that the ‘normal’, but not the ‘tumor’, gene signature, was significantly enriched in E7107-treated LNCaP cells (Supplementary Fig. S11H), suggesting a reversion of LNCaP transcriptome from PCa-like to normal-like. Similarly, pathway analysis in AR^−^p53^−^ PC3 cells identified both convergent (e.g., splicing, differentiation, cell cycle arrest and proliferation inhibition) and unique (i.e., steroid hormone and muscle development) GO categories, when compared to the analysis in LNCaP cells (Fig. 6H). Enrichment of ‘differentiation’ and ‘steroid hormone’ categories in PC3 cells prompted us to examine the androgen/AR signaling. Strikingly, transcript levels of AR itself and many typical AR targets were upregulated in PC3 cells treated with E7107, leading to a dramatic enrichment of AR pathway (Fig. 6I). Furthermore, a LNCaP gene signature was highly enriched in E7107-treated PC3 cells (Fig. 6J). Experimentally, E7107 treatment increased cell size in both PC3 and DU145 cells (Fig. 6K), indicating morphological differentiation. Moreover, although p53 was not activated in PC3 cells due to its genetic loss, several other TS genes (e.g., ALOX15, NKX3-1, RBM4, MIR34A) were upregulated (Fig. 6L). These data, together, suggest a reversal, molecularly and phenotypically, of aggressive PCa cells (PC3) to a more indolent, “LNCaP-like” cell state upon spliceosome inhibition by E7107.

### Therapeutic Targeting of CRPC *in vivo* via Inhibition of Spliceosome Activity

To evaluate the activity of E7107 against CRPC *in vivo*, we treated 3 distinct castration-resistant (AI) PCa xenograft models, i.e., the AR^+/hi^ LNCaP-AI (36), AR^−/lo^ LAPC9-AI (36) and AR^−^ PC3, with E7107 or vehicle (Fig. 7A). The LNCaP-AI and LAPC9-AI models were established by serially passaging the corresponding parent AD tumor cells in castrated immunodeficient mice (36). The LNCaP-AI was initially responsive to Enza but quickly became Enza-resistant whereas LAPC9-AI was refractory to Enza *de novo* (36). Treatment of Enza-refractory LAPC9-AI tumors with either one cycle (i.e., tail vein injection for 5 consecutive days) or two cycles (with 1 week of ‘drug holiday’ between the 2 cycles) effectively inhibited tumor growth (Fig. 7B and 7C; Supplementary Fig. S12A and S12B, left). Similarly, treatment of mice bearing LNCaP-AI with two cycles of E7107 (Fig. 7D; Supplementary Fig. S12C, left) and PC3 xenografts with one cycle of E7107 (Fig. 7E; Supplementary Fig. S12D, left) also inhibited tumor growth. Although a certain degree of toxicity of E7107 was observed, treated mice returned to the range of normal body weight within a week after cessation of treatment (Supplementary Fig. S12A-S12D, right). The endpoint tumors frequently displayed a more ‘differentiated’ morphology manifested by an enrichment of enlarged and polynucleated cells (Supplementary Fig. S12E).

**Figure 7.**
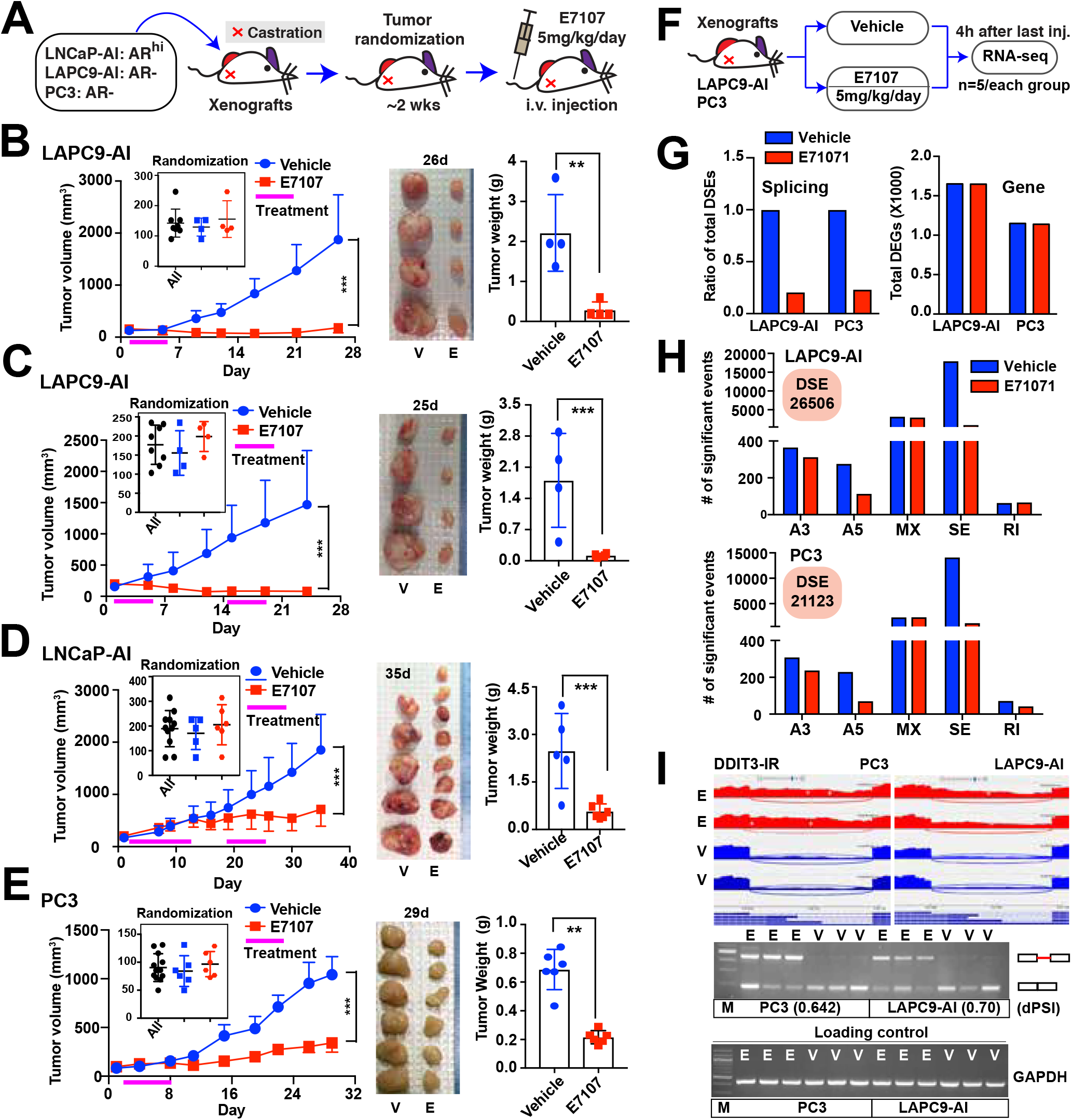
Therapeutic targeting of CRPC cells *in vivo*. (**A**) Schematic of *in vivo* E7107 treatment. (**B-E**) Inhibitory effect of E7107 on the growth of indicated Enza-resistant CRPC models *in vivo*. Shown are growth curve (left), endpoint tumor image (middle) and tumor weight (right) of LAPC9-AI (B and C), LNCaP-AI (D) and PC3 (E) models treated with vehicle or E7107. (**F-G**) Effect of E7107 on CRPC transcriptome *in vivo*. Shown are schematic of RNA-seq experiment (F) and the ratio of total DSEs (G; left) and total DEGs (G; right) identified upon E7107 treatment in indicated CRPC models. (**H**) AS pattern showing that E7107 reshapes the splicing landscape of CRPC xenografts *in vivo*. (**I**) Sashimi plots and RT-PCR validation of IR in DDIT3 gene after E7107 treatment *in vivo*. The event ΔPSI values calculated by rMATS were provided in parentheses. E, E7107; V, vehicle.

To determine whether the tumor-inhibitory effects of E7107 are associated with spliceosome inhibition, we performed RNA-seq analysis in LAPC9-AI and PC3 tumors 4 h after the fifth injection of E7107 (Fig. 7F; see Methods). Consistent with the in vitro data (Fig. 6E), E7107 suppressed the AS globally in both AR^−/lo^ CRPC models (Fig. 7G), evidenced by decreases in A3, A5, and SE (Fig. 7H). Sashimi plot visualization and RT-PCR were performed to validate our splicing analysis (Fig. 7I; Supplementary Fig. S12F). GO analysis of the top 1000 genes bearing down-regulated SE events in LAPC9-AI model upon E7107 treatment revealed an enrichment of several cancer-promoting functional categories including ‘cell cycle and proliferation’, ‘DNA repair’, ‘splicing’, and ‘cancer pathways’ (Supplementary Fig. S12G). Analysis of the gene expression changes after E7107 treatment revealed 3299 and 2289 DEGs, respectively, in LAPC9-AI and PC3 systems without obvious bias on transcription (Fig. 7G). qRT-PCR analysis in tumor samples confirmed the differential expression of selected genes (Supplementary Fig. S13A).

GO analysis of the genes upregulated in E7107-treated LAPC9-AI revealed a broad spectrum of functional categories linked to inhibition of cell proliferation and promotion of normal (prostate) developmental, differentiation, inflammation, and TS pathways, among others (Supplementary Fig. S13B). Since the LAPC9-AI has the AR^−/lo^ phenotype (36), transcription of AR signaling, as expected, remained unaltered (Supplementary Fig. S13C). Notably, gene signatures specific to pri-PCa and CRPC were significantly enriched in E7107-vs. vehicle-treated LAPC9-AI tumors, respectively (Supplementary Fig. S13D), again suggesting that spliceosome inhibition by E7107 reverses the gene expression pattern of LAPC9-AI from CRPC-Ad-like (more aggressive) to pri-PCa-like (less aggressive). We have recently shown that LAPC9-AI molecularly resembles CRPC-Ad (36). In the PC3 model, we observed an increase in the expression of genes involved in muscle development, inflammation, immune cell infiltration, and androgen response, among others, after E7107 treatment (Supplementary Fig. S13E). Interestingly, despite the upregulated category of ‘androgen response’, AR signaling and many targets remained inactivated (Supplementary Fig. S13F). Compared with in vitro data showing that E7107 strongly boosted the AR signaling (Fig. 6I), this discrepancy could be explained by an in vivo environment lacking androgen in castrated hosts such that the ‘E7107 reprogramed’ AR^+^ PC3 cells may not survive. Endpoint PC3 tumors tended to be less aggressive in terms of molecular signatures (Supplementary Fig. S13G-I). For instance, E7107 treatment significantly inhibited pathways associated with cancer metastasis and stemness (Supplementary Fig. S13G), decreased the expression of a PC3-cell signature (Supplementary Fig. S13H), and reverted the gene expression pattern from CRPC-NE like to CRPC-Ad like (Supplementary Fig. S13I). We have previously demonstrated that PC3 cells molecularly resemble the CRPC-NE (7,36).

## DISCUSSION

Studies of AR variants, ARv7 in particular, have implicated splicing dysregulation in PCa resistance to ADT/Enza (20). Recently, splicing factor HNRNPL was identified as a dependency for LNCaP cells (70) and SFPQ (i.e., PSF) was reported to promote AR splicing and CRPC cell survival (71). An examination of race-specific AS changes in PCa in African American (AA) vs. European American (EA) men discovered an AA-enriched PIK3CD isoform that promotes tumor aggressiveness and drug resistance (72). Globally, splicing dysregulation in pri-PCa vs. normal tissues has been observed (13,19). While these studies implicate splicing dysregulation in PCa pathogenesis, the global AS landscape unraveling the dynamic evolution of PCa has not been reported and the impact of aberrant AS alterations on PCa progression, therapy resistance, and patient outcome remains unclear.

Here, we provide the first comprehensively annotated splicing map in PCa using clinical and experimental RNA-seq data covering the entire spectrum of PCa development and progression. Aberrantly spliced genes specific to different PCa stages are both convergently and uniquely enriched in diverse GO terms and pathways linked to many key cellular processes important for cell survival, which establishes aberrant splicing as a distinct mechanism (vs. gene expression regulation) driving PCa progression and therapy resistance. In particular, we observe increasing severity of AS dysregulation and identify IR as a hallmark of stemness and aggressiveness during PCa progression and therapy resistance. Recently, widespread IR, associated with somatic single-nucleotide variations in six cancer types (excluding PCa), has been observed to be more enriched in TS genes leading to their loss of expression (73). Surprisingly, we did not observe a similar trend in PCa. Rather, our data reveals that IR generally enhances gene expression and thus likely functions in PCa biology, suggesting different roles of IR in distinct cancer types. Particularly, we find that IR in PCa impacts genes involved in stemness and cancer-promoting functions (Supplementary Fig. S4G-I), and that AR regulates a splicing program, but not IR specifically, distinct to its transcriptional regulation, suggesting IR as a PCa-regulating mechanism independent of AR axis. In fact, we have observed a general negative association of AR activity with IR level in multiple clinical datasets. Together, our results may establish IR as a common mechanism of cellular stemness, as supported by studies in mouse ESCs (49). The IR prevalent in PCa is not associated strongly with *cis-*genomic features, but seems to be regulated by *trans*-regulatory mechanisms involving the combinatorial effects of multiple SRGs (Fig. 1G and 1H). In support, candidate RBPs modulate not only the IR but also other splicing types as well (Fig. 1H). Alternatively, besides altered spliceosome activity, IR might also be modulated by other molecular alterations. For example, loss-of-function mutations in SETD2 (a H3K36 methyltransferase) and subsequent loss of H3K36 trimethylation at target exons are associated with increased IR in renal cancers (74). Our work expands the view of molecular complexity underlying and justifies further exploration on the role of IR in PCa etiology and progression.

There are many ways by which RNA splicing can be dysregulated in cancer relative to normal cells. Previously, recurrent point mutations in core spliceosome genes (e.g., SF3B1, U2AF1, SRSF2, ZRSR2) have been identified as the driving mechanism underpinning splicing dysregulation in hematological cancers (12). Our genomic analyses of SRGs reveal copy-number variations (CNVs) as the main driver of AS alterations in PCa, which generally alter the expression of affected SRGs and illustrate cancer type-specific differences in mechanisms of splicing dysregulation. Remarkably, the majority of the top altered SRGs are located in regions containing either TS genes or oncogenes (Supplementary Fig. S8A), all of which have not been highlighted in previous large-scale DNA sequencing studies. This raises an interesting question of whether these alterations are just passenger mutations or they causally contribute to PCa pathogenesis. While direct functional evidence implicating them in PCa biology awaits experimental validation, the involvement of these genes in other types of cancer has been reported (9,10,20). Particularly, splicing dysregulation has been recently proposed as a ‘driver’ of transformation independently of oncogenic processes (17). Intriguingly, CNVs of SRGs exerts, comparatively, a dramatic impact on global splicing landscape in CRPC (data not shown), suggesting an enhanced dependency of CRPC on aberrant spliceosome activity. Therefore, these mutated SRGs may bear some of the ‘driver’ properties, and it would be interesting to dissect, in future studies, whether deletion or amplification of CNV-associated SRGs with or without collateral alterations in RB1 or MYC loci, or vice versa, could change cancer phenotypes. Another potential mechanism that may cause splicing abnormality is the mutations in splice sites (12). However, mutations in splice sites constitute the minority of all somatic mutations (as low as ~0.6%) in PCa (75); consequently, we reason that deregulation of SRGs is the main mechanism underpinning splicing abnormalities. In support, the majority of SRGs are mis-expressed in various stages of PCa, consistent with studies showing that altered expression of SRGs, even in the absence of mutations, promotes oncogenesis (10). Of clinical significance, our study has identified many SRGs that can be linked, individually or in combination, to clinical features of advanced PCa, indicating a ‘biomarker’ value. Almost all of these identified prognostic SRGs and DSEs have not previously been implicated in PCa, thus warranting further investigation. Notably, the unfavorable SRG signature that we developed herein predicts PCa progression and correlates with poor patient survival, associates with much ‘twisted’ splicing landscape, and establishes splicing misregulation as a promoter of PCa aggressiveness.

Multiple lines of evidence reveal a preferential dependency of CRPC on aberrant spliceosome activity. *First*, the number of DSEs increases exponentially along the spectrum of cancer progression (Supplementary Fig. S1D), linking the severity of splicing dysregulation to PCa aggressiveness. *Second*, amplifications of SRGs are predominantly observed, and CNVs of SRGs mainly impact global splicing in CRPC. *Third*, more SRGs are dysregulated in CRPC, highlighting a potentially critical role of SRG misexpression in driving CRPC evolution. *Fourth*, the majority of altered SRGs are predictive of worse patient outcome and the unfavorable SRG signature associates with high tumor grade and more prominent disruption in the splicing landscape. *Fifth*, chemical castration and Enza, both of which target AR signaling, reshape the splicing landscape in PCa cells, and the distorted splicing landscape likely contributes to subsequent treatment failure and disease progression (Fig. 2), as documented in other cancer types (12). *Finally*, E7107, the spliceosome modulator, effectively inhibits the growth of multiple experimental CRPC models in vivo (Fig. 6 and 7) regardless of the AR status. In this study, we did not explore the combination of E7107 and Enza because all CRPC models we utilized are already Enza-resistant. A phase-I study of E7107 in patients with advanced solid tumors (PCa excluded) was terminated due to side effects (66) and we also observed certain toxicities of E7101 in animals (this study), suggesting the need to define intricate treatment window and doses for E7101. Our results may point to a new strategy of administering E7107, or other splicing inhibitors, as we show that, interestingly, E7107 promotes PCa cell differentiation and reprograms PCa cells from an ‘androgen-insensitive’ to an ‘androgen-sensitive’ state. We thus envision a potential treatment regimen in which CRPC is first subject to a short-term splicing inhibition (to avoid toxicity and also to reprogram aggressive PCa cells) followed by Enza treatment. Ongoing studies are exploring the value of this sequential treatment protocol. Overall, our findings suggest that there may be a therapeutic window for spliceosome modulators in the treatment of CRPC. Future studies that aim to determine the origins and consequences of aberrant splicing in aggressive PCa could enhance our understanding of disease pathogenesis and aid novel drug development.

## ACKNOWLEDGMENTS

We acknowledge the support of several Shared Resources at the Roswell Park Comprehensive Cancer Center (RPCCC), including Histology, Flow and Image Cytometry and Genomics Resources. This project was supported, in part, by grants from the U.S NIH (R01CA155693, R01CA237027 and R01CA240290) and Department of Defense (W81XWH-14-1-0575 and W81XWH-16-1-0575), and from the CPRIT (RP120380) (all to D.G.T.) and by RPCCC and NCI center grant P30CA016056. S.L. and J.W. were supported, in part, by the NIH grant U24CA232979. D.Z. was supported, in part, by the NIH/NCI grant R21CA218635 and Huazhong Agricultural University (HZAU) Startup Fund for Advanced Talents. We apologize to the colleagues whose work was not cited due to space constraint.

## AUTHOR CONTRIBUTIONS

D.Z. and D.T. conceived and designed the study, interpreted data, and wrote the manuscript;

D.Z. and Y.J. performed most experiments;

D.Z., Q.H., H.C., Y.L., J.W. and S.L. conducted Bioinformatic analysis;

A.T. and J.K. provided assistance in animal experiments;

S.B and P.Z. provided reagents and expertise for the spliceosome modulation experiments;

All authors read and approved the manuscript.

## DECLARATION OF INTERESTS

The authors declare no competing financial interests.

## Notes

The authors declare no potential conflicts of interest.

## REFERENCES

1. Siegel RL, Miller KD, Jemal A. Cancer statistics, 2019. CA: a cancer journal for clinicians 2019 doi 10.3322/caac.21551.

2. Zhang D, Jeter C, Gong S, Tracz A, Lu Y, Shen J, et al. Histone 2B-GFP Label-Retaining Prostate Luminal Cells Possess Progenitor Cell Properties and Are Intrinsically Resistant to Castration. Stem Cell Reports 2018;10(1):228–42 doi 10.1016/j.stemcr.2017.11.016.

3. Zhang D, Zhao S, Li X, Kirk JS, Tang DG. Prostate Luminal Progenitor Cells in Development and Cancer. Trends Cancer 2018;4(11):769–83 doi 10.1016/j.trecan.2018.09.003.

4. Zhang D, Tang DG. “Splice” a way towards neuroendocrine prostate cancer. EBioMedicine 2018;35:12–3 doi 10.1016/j.ebiom.2018.08.037.

5. Beltran H, Prandi D, Mosquera JM, Benelli M, Puca L, Cyrta J, et al. Divergent clonal evolution of castration-resistant neuroendocrine prostate cancer. Nat Med 2016;22(3):298–305 doi 10.1038/nm.4045.

6. Ku SY, Rosario S, Wang Y, Mu P, Seshadri M, Goodrich ZW, et al. Rb1 and Trp53 cooperate to suppress prostate cancer lineage plasticity, metastasis, and antiandrogen resistance. Science 2017;355(6320):78–83 doi 10.1126/science.aah4199.

7. Zhang D, Park D, Zhong Y, Lu Y, Rycaj K, Gong S, et al. Stem cell and neurogenic gene-expression profiles link prostate basal cells to aggressive prostate cancer. Nature communications 2016;7:10798 doi 10.1038/ncomms10798.

8. Tang DG. Understanding cancer stem cell heterogeneity and plasticity. Cell research 2012;22(3):457–72 doi 10.1038/cr.2012.13.

9. Sveen A, Kilpinen S, Ruusulehto A, Lothe RA, Skotheim RI. Aberrant RNA splicing in cancer; expression changes and driver mutations of splicing factor genes. Oncogene 2016;35(19):2413–27 doi 10.1038/onc.2015.318.

10. Zhang J, Manley JL. Misregulation of pre-mRNA alternative splicing in cancer. Cancer discovery 2013;3(11):1228–37 doi 10.1158/2159-8290.CD-13-0253.

11. Park E, Pan Z, Zhang Z, Lin L, Xing Y. The Expanding Landscape of Alternative Splicing Variation in Human Populations. Am J Hum Genet 2018;102(1):11–26 doi 10.1016/j.ajhg.2017.11.002.

12. Lee SC, Abdel-Wahab O. Therapeutic targeting of splicing in cancer. Nat Med 2016;22(9):976–86 doi 10.1038/nm.4165.

13. Kahles A, Lehmann KV, Toussaint NC, Huser M, Stark SG, Sachsenberg T, et al. Comprehensive Analysis of Alternative Splicing Across Tumors from 8,705 Patients. Cancer cell 2018;34(2):211–24 e6 doi 10.1016/j.ccell.2018.07.001.

14. Hsu TY, Simon LM, Neill NJ, Marcotte R, Sayad A, Bland CS, et al. The spliceosome is a therapeutic vulnerability in MYC-driven cancer. Nature 2015;525(7569):384–8 doi 10.1038/nature14985.

15. Sebestyen E, Singh B, Minana B, Pages A, Mateo F, Pujana MA, et al. Large-scale analysis of genome and transcriptome alterations in multiple tumors unveils novel cancer-relevant splicing networks. Genome research 2016;26(6):732–44 doi 10.1101/gr.199935.115.

16. Seiler M, Peng S, Agrawal AA, Palacino J, Teng T, Zhu P, et al. Somatic Mutational Landscape of Splicing Factor Genes and Their Functional Consequences across 33 Cancer Types. Cell reports 2018;23(1):282–96 e4 doi 10.1016/j.celrep.2018.01.088.

17. Climente-Gonzalez H, Porta-Pardo E, Godzik A, Eyras E. The Functional Impact of Alternative Splicing in Cancer. Cell reports 2017;20(9):2215–26 doi 10.1016/j.celrep.2017.08.012.

18. Li Y, Sahni N, Pancsa R, McGrail DJ, Xu J, Hua X, et al. Revealing the Determinants of Widespread Alternative Splicing Perturbation in Cancer. Cell reports 2017;21(3):798–812 doi 10.1016/j.celrep.2017.09.071.

19. Ryan M, Wong WC, Brown R, Akbani R, Su X, Broom B, et al. TCGASpliceSeq a compendium of alternative mRNA splicing in cancer. Nucleic Acids Res 2016;44(D1):D1018–22 doi 10.1093/nar/gkv1288.

20. Paschalis A, Sharp A, Welti JC, Neeb A, Raj GV, Luo J, et al. Alternative splicing in prostate cancer. Nat Rev Clin Oncol 2018;15(11):663–75 doi 10.1038/s41571-018-0085-0.

21. Shen S, Park JW, Lu ZX, Lin L, Henry MD, Wu YN, et al. rMATS: robust and flexible detection of differential alternative splicing from replicate RNA-Seq data. Proceedings of the National Academy of Sciences of the United States of America 2014;111(51):E5593–601 doi 10.1073/pnas.1419161111.

22. Alamancos GP, Pages A, Trincado JL, Bellora N, Eyras E. Leveraging transcript quantification for fast computation of alternative splicing profiles. RNA 2015;21(9):1521–31 doi 10.1261/rna.051557.115.

23. Cancer Genome Atlas Research N. The Molecular Taxonomy of Primary Prostate Cancer. Cell 2015;163(4):1011–25 doi 10.1016/j.cell.2015.10.025.

24. Robinson D, Van Allen EM, Wu YM, Schultz N, Lonigro RJ, Mosquera JM, et al. Integrative clinical genomics of advanced prostate cancer. Cell 2015;161(5):1215–28 doi 10.1016/j.cell.2015.05.001.

25. Sowalsky AG, Xia Z, Wang L, Zhao H, Chen S, Bubley GJ, et al. Whole transcriptome sequencing reveals extensive unspliced mRNA in metastatic castration-resistant prostate cancer. Molecular cancer research: MCR 2015;13(1):98–106 doi 10.1158/1541-7786.MCR-14-0273.

26. Wyatt AW, Mo F, Wang K, McConeghy B, Brahmbhatt S, Jong L, et al. Heterogeneity in the inter-tumor transcriptome of high risk prostate cancer. Genome biology 2014;15(8):426 doi 10.1186/s13059-014-0426-y.

27. Rajan P, Sudbery IM, Villasevil ME, Mui E, Fleming J, Davis M, et al. Next-generation sequencing of advanced prostate cancer treated with androgen-deprivation therapy. European urology 2014;66(1):32–9 doi 10.1016/j.eururo.2013.08.011.

28. Mu P, Zhang Z, Benelli M, Karthaus WR, Hoover E, Chen CC, et al. SOX2 promotes lineage plasticity and antiandrogen resistance in TP53- and RB1-deficient prostate cancer. Science 2017;355(6320):84–8 doi 10.1126/science.aah4307.

29. Zhang Z, Pal S, Bi Y, Tchou J, Davuluri RV. Isoform level expression profiles provide better cancer signatures than gene level expression profiles. Genome Med 2013;5(4):33 doi 10.1186/gm437.

30. Rodriguez-Bravo V, Pippa R, Song WM, Carceles-Cordon M, Dominguez-Andres A, Fujiwara N, et al. Nuclear Pores Promote Lethal Prostate Cancer by Increasing POM121-Driven E2F1, MYC, and AR Nuclear Import. Cell 2018;174(5):1200–15 e20 doi 10.1016/j.cell.2018.07.015.

31. Patro R, Duggal G, Love MI, Irizarry RA, Kingsford C. Salmon provides fast and bias-aware quantification of transcript expression. Nature methods 2017;14(4):417–9 doi 10.1038/nmeth.4197.

32. Yae T, Tsuchihashi K, Ishimoto T, Motohara T, Yoshikawa M, Yoshida GJ, et al. Alternative splicing of CD44 mRNA by ESRP1 enhances lung colonization of metastatic cancer cell. Nature communications 2012;3:883 doi 10.1038/ncomms1892.

33. Lee AR, Li Y, Xie N, Gleave ME, Cox ME, Collins CC, et al. Alternative RNA splicing of the MEAF6 gene facilitates neuroendocrine prostate cancer progression. Oncotarget 2017;8(17):27966–75 doi 10.18632/oncotarget.15854.

34. Dvinge H, Bradley RK. Widespread intron retention diversifies most cancer transcriptomes. Genome Med 2015;7(1):45 doi 10.1186/s13073-015-0168-9.

35. Ben-Porath I, Thomson MW, Carey VJ, Ge R, Bell GW, Regev A, et al. An embryonic stem cell-like gene expression signature in poorly differentiated aggressive human tumors. Nature genetics 2008;40(5):499–507 doi ng.127[pii]10.1038/ng.127.

36. Li Q, Deng Q, Chao HP, Liu X, Lu Y, Lin K, et al. Linking prostate cancer cell AR heterogeneity to distinct castration and enzalutamide responses. Nature communications 2018;9(1):3600 doi 10.1038/s41467-018-06067-7.

37. Zhang D, Tang DG, Rycaj K. Cancer stem cells: Regulation programs, immunological properties and immunotherapy. Semin Cancer Biol 2018 doi 10.1016/j.semcancer.2018.05.001.

38. Qin J, Liu X, Laffin B, Chen X, Choy G, Jeter CR, et al. The PSA(-/lo) prostate cancer cell population harbors self-renewing long-term tumor-propagating cells that resist castration. Cell stem cell 2012;10(5):556–69 doi 10.1016/j.stem.2012.03.009.

39. Rycaj K, Cho EJ, Liu X, Chao HP, Liu B, Li Q, et al. Longitudinal tracking of subpopulation dynamics and molecular changes during LNCaP cell castration and identification of inhibitors that could target the PSA-/lo castration-resistant cells. Oncotarget 2016;7(12):14220–40 doi 10.18632/oncotarget.7303.

40. Choi J, Lee S, Mallard W, Clement K, Tagliazucchi GM, Lim H, et al. A comparison of genetically matched cell lines reveals the equivalence of human iPSCs and ESCs. Nat Biotechnol 2015;33(11):1173–81 doi 10.1038/nbt.3388.

41. Middleton R, Gao D, Thomas A, Singh B, Au A, Wong JJ, et al. IRFinder: assessing the impact of intron retention on mammalian gene expression. Genome biology 2017;18(1):51 doi 10.1186/s13059-017-1184-4.

42. Naro C, Jolly A, Di Persio S, Bielli P, Setterblad N, Alberdi AJ, et al. An Orchestrated Intron Retention Program in Meiosis Controls Timely Usage of Transcripts during Germ Cell Differentiation. Developmental cell 2017;41(1):82–93 e4 doi 10.1016/j.devcel.2017.03.003.

43. Ni T, Yang W, Han M, Zhang Y, Shen T, Nie H, et al. Global intron retention mediated gene regulation during CD4+ T cell activation. Nucleic Acids Res 2016;44(14):6817–29 doi 10.1093/nar/gkw591.

44. Braunschweig U, Barbosa-Morais NL, Pan Q, Nachman EN, Alipanahi B, Gonatopoulos-Pournatzis T, et al. Widespread intron retention in mammals functionally tunes transcriptomes. Genome research 2014;24(11):1774–86 doi 10.1101/gr.177790.114.

45. Lu ZX, Huang Q, Park JW, Shen S, Lin L, Tokheim CJ, et al. Transcriptome-wide landscape of pre-mRNA alternative splicing associated with metastatic colonization. Molecular cancer research: MCR 2015;13(2):305–18 doi 10.1158/1541-7786.MCR-14-0366.

46. Paz I, Kosti I, Ares M, Jr., Cline M, Mandel-Gutfreund Y. RBPmap: a web server for mapping binding sites of RNA-binding proteins. Nucleic Acids Res 2014;42(Web Server issue):W361–7 doi 10.1093/nar/gku406.

47. Ray D, Kazan H, Cook KB, Weirauch MT, Najafabadi HS, Li X, et al. A compendium of RNA-binding motifs for decoding gene regulation. Nature 2013;499(7457):172–7 doi 10.1038/nature12311.

48. Ye J, Liang R, Bai T, Lin Y, Mai R, Wei M, et al. RBM38 plays a tumor-suppressor role via stabilizing the p53-mdm2 loop function in hepatocellular carcinoma. J Exp Clin Cancer Res 2018;37(1):212 doi 10.1186/s13046-018-0852-x.

49. Boutz PL, Bhutkar A, Sharp PA. Detained introns are a novel, widespread class of post-transcriptionally spliced introns. Genes & development 2015;29(1):63–80 doi 10.1101/gad.247361.114.

50. Colombo M, Karousis ED, Bourquin J, Bruggmann R, Muhlemann O. Transcriptome-wide identification of NMD-targeted human mRNAs reveals extensive redundancy between SMG6- and SMG7-mediated degradation pathways. RNA 2017;23(2):189–201 doi 10.1261/rna.059055.116.

51. Kumar A, Coleman I, Morrissey C, Zhang X, True LD, Gulati R, et al. Substantial interindividual and limited intraindividual genomic diversity among tumors from men with metastatic prostate cancer. Nat Med 2016;22(4):369–78 doi 10.1038/nm.4053.

52. Schroeder A, Herrmann A, Cherryholmes G, Kowolik C, Buettner R, Pal S, et al. Loss of androgen receptor expression promotes a stem-like cell phenotype in prostate cancer through STAT3 signaling. Cancer research 2014;74(4):1227–37 doi 10.1158/0008-5472.CAN-13-0594.

53. Liu X, Chen X, Rycaj K, Chao HP, Deng Q, Jeter C, et al. Systematic dissection of phenotypic, functional, and tumorigenic heterogeneity of human prostate cancer cells. Oncotarget 2015;6(27):23959–86.

54. Asangani IA, Dommeti VL, Wang X, Malik R, Cieslik M, Yang R, et al. Therapeutic targeting of BET bromodomain proteins in castration-resistant prostate cancer. Nature 2014;510(7504):278–82 doi 10.1038/nature13229.

55. Olsen JR, Azeem W, Hellem MR, Marvyin K, Hua Y, Qu Y, et al. Context dependent regulatory patterns of the androgen receptor and androgen receptor target genes. BMC Cancer 2016;16:377 doi 10.1186/s12885-016-2453-4.

56. Younis I, Berg M, Kaida D, Dittmar K, Wang C, Dreyfuss G. Rapid-response splicing reporter screens identify differential regulators of constitutive and alternative splicing. Mol Cell Biol 2010;30(7):1718–28 doi 10.1128/MCB.01301-09.

57. Baca SC, Prandi D, Lawrence MS, Mosquera JM, Romanel A, Drier Y, et al. Punctuated evolution of prostate cancer genomes. Cell 2013;153(3):666–77 doi 10.1016/j.cell.2013.03.021.

58. Barbieri CE, Baca SC, Lawrence MS, Demichelis F, Blattner M, Theurillat JP, et al. Exome sequencing identifies recurrent SPOP, FOXA1 and MED12 mutations in prostate cancer. Nature genetics 2012;44(6):685–9 doi 10.1038/ng.2279.

59. Grasso CS, Wu YM, Robinson DR, Cao X, Dhanasekaran SM, Khan AP, et al. The mutational landscape of lethal castration-resistant prostate cancer. Nature 2012;487(7406):239–43 doi nature11125[pii]10.1038/nature11125.

60. Hieronymus H, Schultz N, Gopalan A, Carver BS, Chang MT, Xiao Y, et al. Copy number alteration burden predicts prostate cancer relapse. Proceedings of the National Academy of Sciences of the United States of America 2014;111(30):11139–44 doi 10.1073/pnas.1411446111.

61. Taylor BS, Schultz N, Hieronymus H, Gopalan A, Xiao Y, Carver BS, et al. Integrative genomic profiling of human prostate cancer. Cancer cell 2010;18(1):11–22 doi 10.1016/j.ccr.2010.05.026.

62. Gao J, Aksoy BA, Dogrusoz U, Dresdner G, Gross B, Sumer SO, et al. Integrative analysis of complex cancer genomics and clinical profiles using the cBioPortal. Sci Signal 2013;6(269):pl1 doi 10.1126/scisignal.2004088.

63. Ulz P, Belic J, Graf R, Auer M, Lafer I, Fischereder K, et al. Whole-genome plasma sequencing reveals focal amplifications as a driving force in metastatic prostate cancer. Nature communications 2016;7:12008 doi 10.1038/ncomms12008.

64. McDonald ER 3rd, de Weck A, Schlabach MR, Billy E, Mavrakis KJ, Hoffman GR, et al. Project DRIVE: A Compendium of Cancer Dependencies and Synthetic Lethal Relationships Uncovered by Large-Scale, Deep RNAi Screening. Cell 2017;170(3):577–92 e10 doi 10.1016/j.cell.2017.07.005.

65. Tsherniak A, Vazquez F, Montgomery PG, Weir BA, Kryukov G, Cowley GS, et al. Defining a Cancer Dependency Map. Cell 2017;170(3):564–76 e16 doi 10.1016/j.cell.2017.06.010.

66. Eskens FA, Ramos FJ, Burger H, O’Brien JP, Piera A, de Jonge MJ, et al. Phase I pharmacokinetic and pharmacodynamic study of the first-in-class spliceosome inhibitor E7107 in patients with advanced solid tumors. Clin Cancer Res 2013;19(22):6296–304 doi 10.1158/1078-0432.CCR-13-0485.

67. Pellagatti A, Armstrong RN, Steeples V, Sharma E, Repapi E, Singh S, et al. Impact of spliceosome mutations on RNA splicing in myelodysplasia: dysregulated genes/pathways and clinical associations. Blood 2018;132(12):1225–40 doi 10.1182/blood-2018-04-843771.

68. Wang Y, Chen D, Qian H, Tsai YS, Shao S, Liu Q, et al. The splicing factor RBM4 controls apoptosis, proliferation, and migration to suppress tumor progression. Cancer cell 2014;26(3):374–89 doi 10.1016/j.ccr.2014.07.010.

69. Liu C, Kelnar K, Liu B, Chen X, Calhoun-Davis T, Li H, et al. The microRNA miR-34a inhibits prostate cancer stem cells and metastasis by directly repressing CD44. Nat Med 2011;17(2):211–5 doi 10.1038/nm.2284.

70. Fei T, Chen Y, Xiao T, Li W, Cato L, Zhang P, et al. Genome-wide CRISPR screen identifies HNRNPL as a prostate cancer dependency regulating RNA splicing. Proceedings of the National Academy of Sciences of the United States of America 2017;114(26):E5207–E15 doi 10.1073/pnas.1617467114.

71. Takayama KI, Suzuki T, Fujimura T, Yamada Y, Takahashi S, Homma Y, et al. Dysregulation of spliceosome gene expression in advanced prostate cancer by RNA-binding protein PSF. Proceedings of the National Academy of Sciences of the United States of America 2017;114(39):10461–6 doi 10.1073/pnas.1706076114.

72. Wang BD, Ceniccola K, Hwang S, Andrawis R, Horvath A, Freedman JA, et al. Alternative splicing promotes tumour aggressiveness and drug resistance in African American prostate cancer. Nature communications 2017;8:15921 doi 10.1038/ncomms15921.

73. Jung H, Lee D, Lee J, Park D, Kim YJ, Park WY, et al. Intron retention is a widespread mechanism of tumor-suppressor inactivation. Nature genetics 2015;47(11):1242–8 doi 10.1038/ng.3414.

74. Simon JM, Hacker KE, Singh D, Brannon AR, Parker JS, Weiser M, et al. Variation in chromatin accessibility in human kidney cancer links H3K36 methyltransferase loss with widespread RNA processing defects. Genome research 2014;24(2):241–50 doi 10.1101/gr.158253.113.

75. Kumar A, White TA, MacKenzie AP, Clegg N, Lee C, Dumpit RF, et al. Exome sequencing identifies a spectrum of mutation frequencies in advanced and lethal prostate cancers. Proceedings of the National Academy of Sciences of the United States of America 2011;108(41):17087–92 doi 10.1073/pnas.1108745108.

